# Interleukin-10 knockout mice do not reliably exhibit macroscopic inflammation: a natural history endoscopic surveillance study

**DOI:** 10.1101/2021.12.18.472865

**Authors:** Seung Young Kim, Jaeho Park, Gabriela Leite, Mark Pimentel, Ali Rezaie

**Author notes:** **Correspondence to:** Ali Rezaie, MD, MSc, FRCPC, Medical Director, GI Motility Program, Cedars-Sinai Medical Center, 8730 Alden Drive, Thalians Bldg, E240, Los Angeles, CA, 90048.

## Abstract

**Background:** Interleukin (IL)-10 knockout (KO) mice, used as a model for inflammatory bowel disease (IBD), develop chronic enterocolitis. Endoscopy, the gold standard for evaluation of human mucosal health, is not widely available for murine models. We aimed to assess the natural history of left-sided colitis in IL-10 KO mice via serial endoscopies.

**Methods:** BALB/cJ IL10 KO mice underwent regular endoscopic assessments from 2 up to 8 months of age. Procedures were recorded and blindly evaluated using a 4-component endoscopic score: mucosal wall transparency, intestinal bleeding, focal lesions and perianal lesions (0-3 points each). An endoscopic score ≥1 point was considered as the presence of colitis/flare.

**Results:** IL-10 KO mice (N=40, 9 female) were assessed. Mean age at first endoscopy was 62.5±2.5 days; average number of procedures per mouse was 6.0±1.3. A total of 238 endoscopies were conducted every 24.8±8.3 days, corresponding to 124.1±45.2 days of surveillance per mouse (13.6 years cumulative surveillance). Thirty-three endoscopies in 24 mice (60%) detected colitis, mean endoscopy score 2.5±1.3 (range: 1-6.3). Nineteen mice (47.5%) had one episode of colitis and 5 (12.5%) had 2-3 episodes. All exhibited complete spontaneous healing on subsequent endoscopies.

**Conclusions:** In this largest endoscopic surveillance study of IL-10 KO mice, 40% of mice did not develop endoscopic left-sided colitis. Furthermore, IL-10 KO mice did not exhibit persistent colitis and universally exhibited complete spontaneous healing without treatment. The natural history of colitis in IL-10 KO mice may not be comparable with that of IBD in humans and requires careful consideration.

## Introduction

Interleukin-10 (IL-10) is an immunoregulatory cytokine that is essential for the maintenance of intestinal homeostasis, and is secreted by a wide range of immune cells including macrophages, dendritic cells, and T cells.(8) IL-10 targets both innate and adaptive immune responses and helps to preserve tissue integrity and promote tissue homeostasis through its anti-inflammatory, anti-apoptotic, and tissue-regenerating properties.(12)

The IL-10 knockout (KO) mouse is a well-studied model of inflammatory bowel disease (IBD) that develops spontaneous Crohn’s disease-like intestinal inflammation, which is mediated by a Th-1 type cytokine profile and requires an intestinal bacterial microbiota to develop.(14) Colitis in IL-10-deficient mice is characterized by histological findings similar to those in human IBD.(2, 10) The discontinuous and transmural inflammatory lesions are characterized by inflammatory cell infiltrates into the lamina propria and submucosa, epithelial hyperplasia, mucin depletion, crypt abscess, ulcers, and thickening of the intestinal wall.(1, 2)

In this murine colitis model, the severity of intestinal inflammation is mainly determined by histological examination. However, longitudinal assessment via histological examination is not possible as it requires colon harvest after euthanasia. Endoscopic examination in mice has proven a useful research tool and provides for objective assessment of colonic inflammation as well as repeated evaluations of individual animals over time.(7, 9) We have shown that murine colonoscopy has substantial interobserver reliability.(13) Therefore, direct serial examination with endoscopy represents a useful tool to elucidate the natural history of colitis in the IL-10 KO mouse model. In this study, we assessed the course of the development of left-sided colitis in BALB/cJ IL10 KO mice via serial endoscopies.

## Materials and Methods

### Animals

Male and female homozygous IL-10^−/-^ mice BALB/cJ mice (C.129P2(B6)-Il10<tm1Cgn>/J, Jackson Laboratories, Bar Harbor, ME) were housed in a vivarium in an accredited animal facility. Animals are maintained under barrier-protected conditions and bedding is changed weekly. To ensure a homogeneous microbial environment, dirty cage bedding was transferred between cages. Mice consumed a standard diet and water *ad libitum*. The study was approved by the Cedars-Sinai Institutional Animal Care and Use Committee (IACUC #7304).

### Endoscopic examination

Mice underwent regular endoscopic assessments from 2 months of age to up to 8 months of age. No bowel preparation was used prior to endoscopies. An isoflurane anesthetic gas (1-5%) mixture (carrier gas: compressed oxygen) was used to induce anesthesia in a chamber, and once the respiratory rate had slowed to approximately one breath per second, the animals were removed from the induction chamber and maintained under sedation using a nose cone anesthesia. A rigid pediatric cystoscope (Olympus A37027A) was used to assess the intestinal mucosa up to the splenic flexure. All endoscopies were recorded and interpreted blindly by two gastroenterologists (SYK, JHP) with expertise in animal model endoscopies.

Endoscopies were scored using a mouse endoscopic scoring system devised by Kodani et al. (9) (Supplemental Table 1). In this scoring system, two major components are used: 1) assessment of the extent and severity of colorectal inflammation (based on perianal findings, transparency of the wall, mucosal bleeding, and focal lesions), and 2) numerical sorting of clinical cases by their pathological and research relevance based on decimal units with assigned categories of observed lesions and endoscopic complications (decimal identifiers). Score of each component of colorectal inflammation varied from 0 to 3 points. An endoscopic score ≥1 point was considered to indicate the presence of colitis/flare.

## Results

### Endoscopic examination of IL-10 KO mice

A total of 40 IL-10 KO mice (9 female) were assessed via serial endoscopies. Mean age at the time of first endoscopy was 62.5±2.5 days and the average number of endoscopic examinations per animal was 6.0±1.3. A total of 238 endoscopies were conducted every 24.8±8.3 days, cumulatively corresponding to 4,964 days (13.6 years) of endoscopic surveillance. During surveillance periods (124.1±45.2 days per mouse), 16 mice never developed colitis within reach of the endoscope (40%), and 24 mice (60%) exhibited at least one episode of colitis (colitis score ≥1) with a mean endoscopy score of 2.5±1.3 (range: 1-6.3).

The most common finding was erosion or erythema. Mean age at first episode of colitis was 83.7±37.5 days. The prevalence of endoscopic colitis appeared to decrease with age as 22.7% (20/88) of endoscopies exhibited colitis in mice less than 80 days old, as compared to 10.1% (7/69) in mice between 81 and 120 days old, and 7.6% (6/79) in mice older than 120 days.

### Pattern of colitis in IL-10 KO mice

Nineteen mice (47.5%) showed a single flare and 5 mice (12.5%) showed 2 or 3 flares. The colitis pattern was similar between males and females with 45.2% (14/31) and 55.6% (5/9) having a single colitis episode and 12.9% (4/31) and 11.1% (1/9) having multiple episodes of colitis, respectively. There was no difference in distribution of colitis pattern between the two groups. All mice with colitis exhibited complete spontaneous healing on subsequent endoscopies (Figure 1). Three distinct endoscopic phenotypes were observed among IL-10 KO mice (Figure 2). No mice exhibited perianal disease.

**Figure 1.**
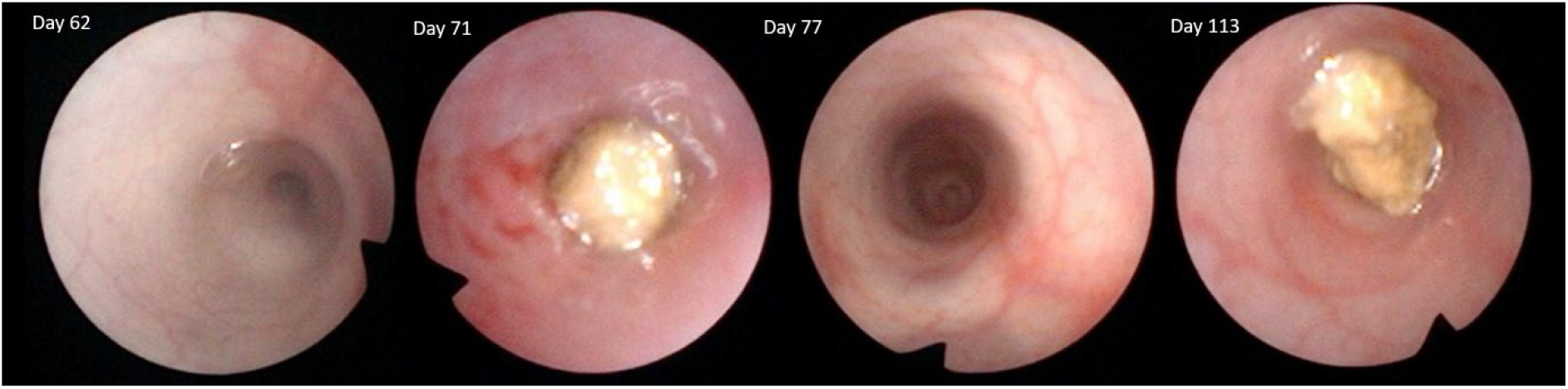
Successive endoscopic images of a mouse colon (mouse age is shown in days). Erosions, loss of vascular markings and mucosal bleeding/friability on day 71 were found to have spontaneously resolved without treatment or intervention on follow-up endoscopies.

**Figure 2.**
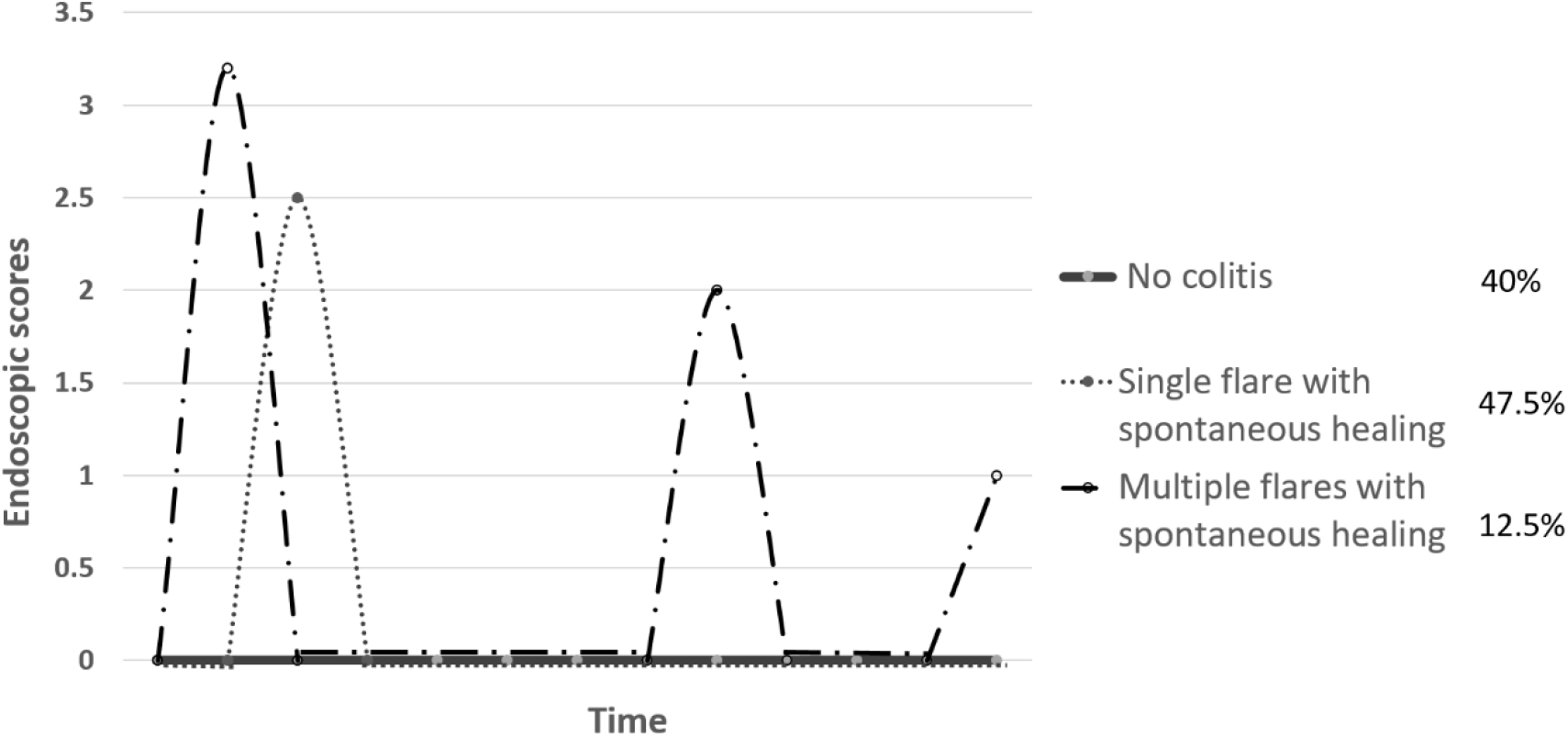
Schematic graph showing three distinct phenotypes of BALB/cJ IL-10 KO mice during endoscopic surveillance.

## Discussion

Repeated blind endoscopic assessments of 40 BALB/cJ IL-10 KO mice, which corresponded to a total of 13.9 years of surveillance, revealed that only 60% of mice developed transient left-sided colitis with a variable and unpredictable recurrence pattern. However, the most striking finding of our study is the universal spontaneous healing of colitis in IL-10 KO mice in the absence of any treatment. To our knowledge, this phenomenon has not been previously reported. Spontaneous resolution of extensive colitis in humans is an unlikely scenario (6); hence, the use of IL-10 KO mice could potentially introduce a challenge in the extrapolation of benchtop results to human subjects. Moreover, lack of persistent endoscopic colitis could make evaluation of drugs difficult, by over- or underestimating therapeutic effects in controlled animal studies. Thus, a potentially beneficial modality could be inadvertently deemed ineffective or vice-versa.

While germ-free IL-10-deficient mice exhibit no histologically detectable colitis(14), IL-10-deficient mice raised under conventional conditions exhibit weight loss, anemia and colitis at the age of 4-8 weeks (10) demonstrating the importance of gut microbiome in development of inflammation in this model. In another study, the rate of spontaneous histologic colitis in IL-10 KO mice (C57BL/6 × 129 Ola) until the age of 12 weeks was nearly 100%(1). In contrast, we found that only 60% of mice developed transient left-sided colitis. This difference could be due to a disconnect in the endoscopic presentation of IL10 KO mice in relation to presence of mural inflammation. In contrast, outright disconnect between histology and endoscopic findings in uncommon in human IBD. The genetic background of our mice may also play a role in low incidence rate of colitis. It has been shown that the genetic background of IL-10 KO mice correlates with susceptibility to colitis and also with the severity of colitis in this model.(8)

The IL-10 deficient mouse can be described as a multi-hit model where a colitogenic trigger initiates the inflammatory process, strain-specific genetic factors determine the dysregulation of the mucosal immune response, and the gut microbiota modify these susceptibilities and responses.(8) Individual IL-10 KO mice in the same colony and even in the same cage develop spontaneous colitis at a non-uniform rate(5). IL-10-deficient C3H mice from the same parental breeding stocks but maintained in two different facilities had significant differences in histopathological scores at the same age. These differences have been attributed to differences in dietary sources and ingredients, and to the water consumed (autoclaved or not).(11) To avoid these potential confounding effects, mice in this study were housed under the same barrier-protected conditions and bedding was changed weekly. Furthermore, all mice were fed the same diet and to ensure a homogeneous microbial environment, dirty cage bedding was transferred between cages.

Our study has several limitations. We did not measure inflammatory markers in stool or blood samples or perform histologic assessments. We also did not take biopsy samples during endoscopies, due high risk of perforation. In clinical practice, the severity of IBD is usually determined through a combination of endoscopic, histologic and radiological findings, and serologic or fecal biomarkers are often used as noninvasive and inexpensive supporting methods. Along with C-reactive protein and erythrocyte sedimentation rate, fecal calprotectin and fecal lactoferrin have become part of the current battery of laboratory tests performed during the clinical management of IBD.(4) Fecal Lipocalin-2 has been reported to serve as a potential biomarker of intestinal inflammation in dextran sulfate sodium-induced colitis and IL-10 KO mouse models.(3) Further longitudinal studies on animal models of colitis are warranted to correlate endoscopic findings with inflammatory markers and histology.

In conclusion, in the largest endoscopic surveillance study of IL-10 KO mice to date, only 60% of mice developed transient left-sided colitis with a variable recurrence pattern. This natural history of colitis in IL10 KO mice may not be comparable to human IBD, and requires careful consideration when drawing conclusions in translational research. Other strains of IL10 KO mice should be assessed to further delineate the natural history of colitis.

## Disclosures

### Authors’ contributions

MP and AR conceived and designed the study. SYK, JHP, GL, and AR performed the experiments. GL, AR and MP analyzed the data. SYK, JHP, GL, and AR wrote the manuscript, and AR and MP revised and edited the manuscript. All authors read and approved the final manuscript.

