## Supplemental Table 1 for "Interleukin-10 knockout mice do not reliably exhibit macroscopic inflammation: a natural history endoscopic surveillance study"

Supplemental Table 1. Endoscopic scoring system for assessment of colitis in mice (adapted from Kodani et al (9))

| Inflammation score | |  |
| --- | --- | --- |
| Wall transparency | | 0 = Yes, small/large vessels are visible. |
|  |  | 1 = Yes; however, most small vessels cannot be seen. |
|  |  | 2 = Not sure; Only very large vessels are seen. |
|  |  | 3 = Not at all; Thickened appearance of mucosa. Blood vessels cannot be seen. |
| Intestinal bleeding | | 0 = No |
|  |  | 1 = Yes; Contact bleeding due to endoscopic trauma. |
|  |  | 3 = Yes; Spontaneous bleeding, not due to endoscopic trauma. |
| Focal lesions | | 0 = No |
|  |  | 1 = Edematous areas of mucosa |
|  |  | 2 = Erosion/reddened area(s) |
|  |  | 3 = Ulcers (often with white material)/Stricture |
| Perianal findings | | 0 = No |
|  |  | 1 = Diarrhea/fecal clumps |
|  |  | 2 = Bloody anal discharge |
|  |  | 3 = Rectal prolapse. Granulation. Fistula |
| Decimal identifiers | |  |
|  | 0.0 - no specific findings | |
|  | 0.1 - perianal fistula | |
|  | 0.2 - erosion | |
|  | 0.3 - ulceration | |
|  | 0.4 - intestinal stricture | |
